# pH-Dependent Silica Nanoshell Degradation Influences SERRS Enhancement in Biological Environments

**DOI:** 10.64898/2026.02.24.707739

**Authors:** William H. Skinner, Samuel Park, Fay Nicolson

**Affiliations:** Department of Radiation Oncology, Dana-Farber Cancer Institute and Harvard Medical School, Boston, MA, 02215, USA

## Abstract

Silica-encapsulated gold nanostars (AuNStar-SiO_2_) are a widely used plasmonic nanoparticle platform for surface-enhanced resonance Raman scattering (SERRS) bio-applications. In this paper, we demonstrate that coupled nanostar subpopulations can dominate the ensemble-average SERRS response of the suspension and that near-neutral standard cell culture conditions are sufficient to hydrolyze the silica nanoshell and introduce variability in signal intensity following *in vitro* endocytosis. Monomeric and oligomeric AuNStar-SiO_2_ fractions were isolated using continuous density-gradient centrifugation and monomeric populations were found to exhibit significantly weaker SERRS compared to their oligomeric counterparts. Using monomer-enriched AuNStar-SiO_2_, we investigated the stability of the silica nanoshell under conditions representative of sequential acidification during endocytosis and characterized the subsequent changes to nanoparticle optical properties. In acidic environments, reflecting lysosomal pH, the silica shell was stable, whereas near-neutral and alkaline conditions in cell culture medium induced silica-shell hydrolysis, nanostar release, and interparticle aggregation, leading to transient SERS amplification. When cells were treated with AuNStar-SiO_2_ under near-neutral and acidic conditions, we observed the opposite trend in SERS signal strength. At pH 7.4, the SERRS signal was suppressed even though transmission electron microscopy (TEM) images of intracellular nanoparticles showed progressive extents of silica hydrolysis, while at pH 6.4 SERS signal was strong and the silica shell of intracellular nanoparticles remained intact. Together, these findings show how SERRS output can differ between control conditions and biological applications, highlighting the role that local environmental factors play in nanoparticle stability and performance. Our results highlight the previously overlooked role of silica nanoshell instability on SERRS signal output in physiological environments and describe opportunities to harness silica nanoshell hydrolysis to improve the biomedical application of silica-coated plasmonic probes.

## INTRODUCTION

Plasmonic gold nanoparticles have emerged as versatile tools for biological applications due to their biocompatibility and optical properties.^1^ The excitation of localized surface plasmon resonances (LSPR) in these particles generates strong field confinement at their surfaces, facilitating surface-enhanced Raman scattering (SERS).^2^ Surface-enhanced resonance Raman spectroscopy (SERRS) further amplifies Raman scattering signals by coupling the intrinsic electronic transitions of reporter dye molecules with the electromagnetic enhancement produced in the near-field of plasmonic nanostructures.^3^ SE(R)RS has been applied *in vitro* and *in vivo*, where bright, spectrally distinct signals enable nanoparticle-based imaging. *In vitro*, SE(R)S has been applied to image nanoparticle endocytosis in adherent cancer cell lines,^4^ quantify nanoparticle binding to cancer cell membrane receptors,^5^ and image cells in three-dimensional tissue structures.^6,7^ *In vivo*, SE(R)RS nanoparticles have been used as contrast agents for noninvasive tumor mapping and to support efficient tumor resection.^8–14^ The nanoscale dimensions of these materials enable cellular internalization via endocytosis and the opportunity to measure intracellular physiological changes such as pH,^15^ reactive oxygen species generation,^16,17^ miRNA production,^18^ and temperature.^19^

The electromagnetic contribution to SER(R)S arises from strong local field enhancements occurring when the LSPR of metallic nanostructures is excited. The most significant enhancements are typically localized at high-curvature features, such as the tips of nanorods or nanostars.^20,21^ Significant enhancement can also occur within nanogaps formed between adjacent plasmonic particles, where plasmonic hybridization produces confined “hot-spot” regions.^22–24^ In practice, both electromagnetic enhancement mechanisms can coexist within a heterogeneous sample, and their relative contributions are rarely characterized in application-focused SE(R)RS studies. Because shifts between the two regimes can markedly alter SE(R)RS intensity, characterizing the relative contributions of these effects is essential for a full understanding of SERS outputs in sensing and imaging experiments.

Purification of heterogeneous nanoparticle systems into monomeric, dimeric, and higher-order assemblies has been identified as a critical step toward harnessing the full potential of plasmonic nanostructures, owing to the new optical properties that develop when plasmonic nanoparticles couple.^25–27^ Multiple studies have demonstrated that density-gradient centrifugation can effectively separate subpopulations of coupled nanospheres into fractions comprising dimers, trimers, and tetramers that exhibit markedly stronger SERS than monomers, indicating that monomeric nanospheres contribute minimally to the overall measured signal. Therefore, isolating and characterizing the subpopulations that drive SE(R)RS enhancement is essential for improving the translational potential of these nanoparticles for imaging and sensing.

SE(R)RS-active nanoparticles are readily endocytosed by many mammalian cell lines, enabling their application as intracellular sensing and imaging agents.^15,28,29^ Following endocytosis, nanospheres are concentrated within vesicles and ultimately trafficked to lysosomes, creating electromagnetic hotspots between particles and enhancing intracellular SERS.^28,30,31^ The localization of coupled nanoparticles within the lysosome has enabled label-free spectral analysis of protein corona degradation by the acidic and enzyme-rich lysosomal environment^31^ and the analysis of lipid accumulation in lysosomes following drug treatment.^32^ Long-term studies have shown that gold nanoparticles persist in the lysosomes of liver cells for up to 7 weeks, with transmission electron microscopy (TEM) images showing retention of nanoparticle morphology for both nanospheres and rods.^33^ However, the assumption that gold nanoparticles are biologically inert in lysosomes has been challenged by TEM studies in fibroblasts, which show that gold nanoparticles are degraded by reactive oxygen species and recrystallize into 2.5 nm crystalline gold particles within the lysosome.^34^ Together, these studies demonstrate the potential of SE(R)S for intracellular sensing, while underscoring that both the nanoparticle surface coatings and the nanoparticles themselves can undergo significant chemical degradation following endocytosis.

Silicated gold nanoparticles have demonstrated promise as *in vitro* SERRS tags^5,35^ and as *in vivo* contrast agents.^8,9,11,36,37^ However, the origin of the SERRS signal and the role of strongly emitting subpopulations of particles are rarely characterized in these platforms prior to application. In the present study, we apply centrifugal purification to AuNStar-SiO_2_ nanoparticles to identify and characterize the dominant subpopulations responsible for SERRS enhancement. Using these well-defined fractions, we then examine how silica shell stability and the associated SERRS signal evolve under conditions that mimic the progressively acidic environments encountered during endocytic trafficking *in vitro*. We show that silica shell dissolution can both enhance or suppress the SERRS signal depending on the spectral acquisition time point and the chemical environment. Finally, we use TEM to investigate silica shell stability following endocytosis of the nanoparticles, probing how the transition from the near-neutral extracellular environment to the acidic intracellular environment alters nanoparticle structure and SERRS spectral intensity in human colorectal adenocarcinoma cell line SW48.

## MATERIALS AND METHODS

### Materials

Gold(III) chloride trihydrate (Sigma Aldrich, #520918), tetraethyl orthosilicate (Sigma Aldrich, #333859), sodium borohydride (Sigma Aldrich, #213462), N,N-dimethylformamide (Sigma Aldrich, #227056), glycerol (Sigma Aldrich, #65516), ammonium hydroxide (Sigma Aldrich #221228), IR780p (BOCSci, CAS: 23178-67-8), IR-792 (Sigma Aldrich, CAS: 207399-10-8), IR-780I (Sigma Aldrich, CAS: 207399-07-3), IR820 (Sigma Aldrich, CAS:172616-80-7), IR-140 (Sigmal Aldrich, CAS:53655-17-7), IR-775 (Sigma Aldrich, CAS: 199444-11-6), IR-792p (Sigma Aldrich, CAS: 207399-10-8), ethanol (Sigma Aldrich, #459844), phosphate-buffered saline (Corning, 21-040-CV), and *iso*-Propyl Alcohol (Sigma Aldrich, PX1838-1). Glutaraldehyde, 2 % paraformaldehyde, 2% Tannic acid, osmium tetroxide, and uranyl acetate (Electron Microscopy Sciences). 0.1 M Sodium Cacodylate (Sigma Aldrich), LX112 resin (Ladd Research Industries, Williston, VT). All chemicals were used as received unless otherwise stated. Ultrapure water 18 MΩ⋅cm, was used in all experiments.

### AuNStar synthesis

AuNStar’s were fabricated following published methods with minor modifications.^11,38,39^ 5 nm gold seeds were synthesized by adding 2 mL of 25 mM HAuCl_4_ solution to 200 mL of MilliQ purified water, followed by addition of freshly prepared 6 mL of ice-cold 100 mM NaBH_4_ solution under vigorous stirring. After 30s, the resulting mixture was diluted fivefold with MilliQ purified water and left to stand overnight at room temperature. Bare gold nanostars were synthesized by dissolving 3.5 g of ascorbic acid in 100 ml of ultrapure water at 4°C under rapid stirring and adding 380 µl of gold seed followed by 110 µl of 400 mM HAuCl_4_. The solution was stirred for 3 minutes at 4 °C and centrifuged at 4200 rcf at 4 °C for 20 minutes. The supernatant was discarded, and the remaining 1 ml AuNStars were transferred to a dialysis cassette (3,500 MWCO, Thermo Scientific) and dialyzed against ultrapure water for three days at 4 °C with daily water changes.

### IR dye SERS test on bare AuNStars

The NIR dyes IR820, IR783, IR792, IR780p, IR780I, IR140, and IR775 were purchased as is and dissolved in dimethylformamide at 20 mM. AuNStars were diluted to an LSPR extinction of 0.3 in water and EtOH. IR dyes were added to a final concentration of 7 nM for IR783, IR780p, IR140, and IR775, 2 nM for IR792 and IR780I, and 3 nM for IR820. Solutions were vortexed, and SERS was measured immediately.

### AuNStar silication with IR780p dye

Silication was performed following published methods with minor adjustments.^9,11,36,39,40^ Silication reagents were initially added to separate falcon tubes, A and B. 25 ml of isopropanol, 1 ml of tetraethyl orthosilicate (TEOS), 100 μl of 20 mM IR780p in dimethylformamide, and 2.5 ml of ultrapure water were added to tube A. 10 ml of ethanol, 1 ml of AuNStars and 500 μl of NH_4_OH, were added to tube B. Tube B was added to tube A under rapid vertexing and the mixture was placed on a shaker for 20 minutes at 300 rpm before quenching with 60 ml ethanol and centrifugation at 3200 rcf. The resulting pellet was sonicated and washed three times with ethanol, centrifuging at 10,000 rcf. The AuNStar-SiO_2_ were then washed with water once and resuspended in 100 µl of ultrapure water.

### Continuous density gradient centrifugation

A continuous density gradient was generated using glycerol in water mixtures. Solutions of 30%, 35%, 40%, 45%, and 50% glycerol in water (v/v) were prepared, and 2 mL of each solution was carefully added to a 15 mL centrifuge tube, starting with the 30% solution and sequentially increasing the glycerol content to 50%.^26^ The tube was then carefully placed horizontally to the bench for 5 minutes, after which it was centrifuged for 5 minutes at 3200 rcf to facilitate the formation of the continuous density gradient. 100 µl of AuNStar-SiO_2_ were then carefully added to the top of the continuous density gradient and centrifuged for 20 minutes at 3200 rcf. Samples were collected from the top of the density gradient using a 1000 µL pipette and divided into seven fractions based on extraction depth (Figure 3A). Fraction 1 was the uppermost 0-1.5 mL of the density gradient. Fractions 2-7 were subsequently collected in 1 mL increments corresponding to 1.5-2.5 mL (fraction 2), 2.5-3.5 mL (fraction 3), 3.5-4.5 mL (fraction 4), 4.5-5.5 mL (fraction 5), 5.5-6.5 mL (fraction 6), and 6.5-7.5 mL (fraction 7). The remaining volume of the density gradient was discarded. Each fraction was washed three times with ultrapure water, and pelleting was performed with centrifugation at 10,000 rcf for 5 minutes.

### Nanoparticle characterization techniques

All samples were diluted in ultrapure water to an LSPR extinction of 0.3 for characterization. Extinction spectra were collected with a UV-26000 UV-Vis Spectrophotometer (Shimadzu). DLS and zeta potential measurements were taken on a Zetasizer Nano series (Malvern Panalytical). Parameters for Raman measurements are outlined in the “Raman Measurements” section. For TEM, AuNStar-SiO_2_ were suspended in ethanol, and 10 μl was dried on TEM grids before imaging with a JEOL JEM 1400 Plus microscope.

### Silica shell hydrolysis

Phosphate-buffered saline (PBS) was confirmed to have a pH of 7.4 using a pH meter and then adjusted to pH 4, 6.5, and 9 by adding 0.1 M HCl or 0.1 M NaOH. AuNStar-SiO_2_ were purified by continuous density gradient centrifugation, and fraction 3 was resuspended at an extinction of 0.3 at the LSPR in pH 4, 6.5, 7.4, and 9 PBS. SERS, UV-vis, and DLS measurements were taken at regular intervals for 25 h. Samples were vortexed before they were measured and stored at 37 °C between measurements.

### *In vitro* treatment with AuNStar-SiO_2_

Human colorectal adenocarcinoma (SW48) cell line authenticity was confirmed by short tandem repeat (STR) testing, and the cell line was tested for mycoplasma. Cells were cultured in 60 mm Petri dishes in RPMI 1640 medium (Gibco) containing 10% fetal bovine serum (FBS) and incubated at 37 °C in 5% CO_2_. Cells were passaged with TrypLE Express (Gibco) up to 20 passages. For Raman experiments, cells were seeded at 200,000 cells per 35 mm plate and cultured for three days. Six plates were prepared for each experiment. On the day of AuNStar-SiO_2_ treatment, the cells were washed with PBS (×3) and 2 ml of phenol-free RPMI (0% FBS). Phenol-free RPMI (0% FBS) was pre-incubated for 30 minutes at 37 °C in 5% CO_2_ to ensure complete CO_2_/bicarbonate buffering. The RPMI was adjusted to pH 6.4 by adding 20 µl/ml of 1 M HCl, and the pH was confirmed with a pH meter. RPMI at pH 7.4 was used as is and confirmed with a pH meter. Then, 2 ml of pH 7.4 or pH 6.4 phenol-free RPMI (0% FBS), was mixed with fraction 3 AuNStar-SiO_2_ to an LSPR extinction of 0.3, and immediately added to the cells. Three petri dishes were treated with each pH condition. The cells were incubated with AuNStar-SiO_2_ for 4 hours at 5% CO_2_ and 37 °C, then washed with PBS (×3). For fixed-cell Raman mapping experiments, the cells were treated with 4% paraformaldehyde for 10 minutes, followed by PBS washes (×3) and submersion in 2 ml of ultrapure water for mapping. For live-cell Raman experiments, the cells were dissociated with TrypLE™ and pelleted at 500 × g for 10 minutes, and the supernatant was completely removed before Raman spectral acquisition.

### Raman measurements

All Raman measurements from bare AuNStar and AuNStar-SiO_2_ in solution were collected using a 785 nm laser (Innovative Photonics Solutions) coupled to a fiber optic probe (Innovative Photonics Solutions) and a high-throughput f/2 spectrometer (Innovative Photonics Solutions). For each measurement, 1 ml of nanoparticle suspension was placed in a plastic cuvette and inserted into a custom-built cuvette holder; all cuvette measurements were collected at 100 ms exposure and 40 mW power. For live-cell Raman measurements, SW48 cell pellets treated with AuNStar-SiO_2_ were transferred onto a glass slide, and spectra were acquired by focusing a fiber-optic Raman probe (Innovative Photonic Solutions) directly onto the pellet using a 1 ms exposure time and a laser power of 40 mW. Raman maps of fixed cells were acquired with the XploRA confocal microscope (HORIBA) using a 10 × objective, 10% laser power, 0.1 s acquisition, and a step size of 4 µm across a 320 × 530 µm area. Three maps from different regions of the petri dish were collected per sample. In both SW48 cell experiments, three replicates were analyzed per pH condition.

### TEM of intracellular AuNStar-SiO_2_

SW48 cells were seeded on a glass cover slip in a 35 mm petri dish and treated with AuNStar-SiO_2_ as outlined in the section “*In vitro* treatment with AuNStar-SiO_2_”. Following PBS washes, cells were fixed in 2.5% glutaraldehyde (diluted from 25% glutaraldehyde (aq)), 2% paraformaldehyde (diluted from 16% aqueous paraformaldehyde), 2% Tannic acid (w/v) in 0.1 M Sodium Cacodylate pH 7.4 for 1 h at room temperature. Cells were washed with 0.1 M Sodium Cacodylate, and then post-fixed for 1 h at 4 °C in 1% osmium tetroxide in 0.1 M Sodium Cacodylate. Cells were then washed in deionised water and incubated in 2% aqueous uranyl acetate overnight at 4 °C. The following day, tissues were washed with deionised water and then dehydrated at 4 °C in a graded ethanol series. Tissues were then brought to room temperature and dehydrated with 100% ethanol. Infiltration with LX112 resin, was followed by embedding in flat bottom Beem capsules cut to form a cylindrical mold on the growing substrate. The resulting blocks were separated from the culture plates using liquid nitrogen and then sectioned using a Leica Ultracut E ultramicrotome and sections placed on formvar and carbon-coated grids. The sections were contrast-stained with 2% uranyl acetate followed by lead citrate. The samples were imaged with a JEOL JEM 1400 Plus microscope.

### Data processing

All spectra collected with the fiber optic probe were baselined using adaptive iteratively reweighted Penalized Least Squares (airPLS) in MATLAB (R2024b).^41^ In the case of assessment of SERRS signal from gold nanostars and IR dyes, spectra were collected from SERRS-active samples contained in a plastic cuvette. Background contributions from the plastic were removed by scaling and subtracting a Raman spectrum of a water-filled cuvette. Scaled background subtraction was also performed to remove the ethanol contribution from spectra acquired when AuNStars were dispersed in ethanol. In experiments with replicates, the mean was calculated, and the error was computed using the sample standard deviation and propagated via the root-sum-square formula. Raman spectra collected from the cell pellet were baseline-corrected using airPLS, and the mean (±SD) spectral intensity at each wavenumber was calculated. For mapping experiments, spectra were baselined in LabSpec6 (HORIBA, version 6.8.1.9) and exported to MATLAB. Baseline noise was calculated between 815 cm^-1^ and 963 cm^-1^ from spectra without SERS signal. The signal intensity at 940 cm^-1^, the SERS peak of AuNStar-SiO_2_ was calculated for each pixel.^42,43^ Pixels with a signal-to-noise (SNR) ratio of greater than 3 were assigned a value of 1 in the matrix (i.e. hot pixels), and pixels with an SNR less than 3 were assigned a value of 0. A mask was drawn around cells in the bright-field image and applied to exclude pixels outside the cell area in the matrix. Hot pixels per µm^2^ of cell coverage were computed from three Raman maps per sample and then averaged to obtain a sample mean. GraphPad Prism (version 10.5.0) was used for statistics and graphing. TEM images were analyzed in ImageJ (version 1.54g). Nanoparticle monomers and oligomers in each fraction were counted manually (>100 particles per fraction from three separate AuNStar-SiO_2_ batches), and circularity measurements were conducted by manually highlighting the periphery of the AuNStars. The size of intracellular nanoparticles was measured manually using the ImageJ length function, with 440 intracellular AuNStar-SiO_2_ particles analyzed in cells treated at each pH.

## RESULTS

### IR dye selection and AuNStar-SiO_2_ synthesis

A wide variety of near-infrared (NIR) dyes have been explored for SERRS imaging applications owing to their strong absorption cross-sections and compatibility with biological imaging windows in the NIR. To identify the most suitable Raman reporter for AuNStars, we screened seven commercially available cyanine dyes reported in the literature.^36,44^ Each dye was added to AuNStars suspended in water or ethanol, and the resulting SERS spectra were recorded (**Figure 1A-G**). Ethanol was selected as a solvent to emulate the chemical environment during the Stöber silication process, which is widely used to encapsulate IR dyes near the AuNStar surface for imaging applications.^9,39,40^

**Figure 1.**
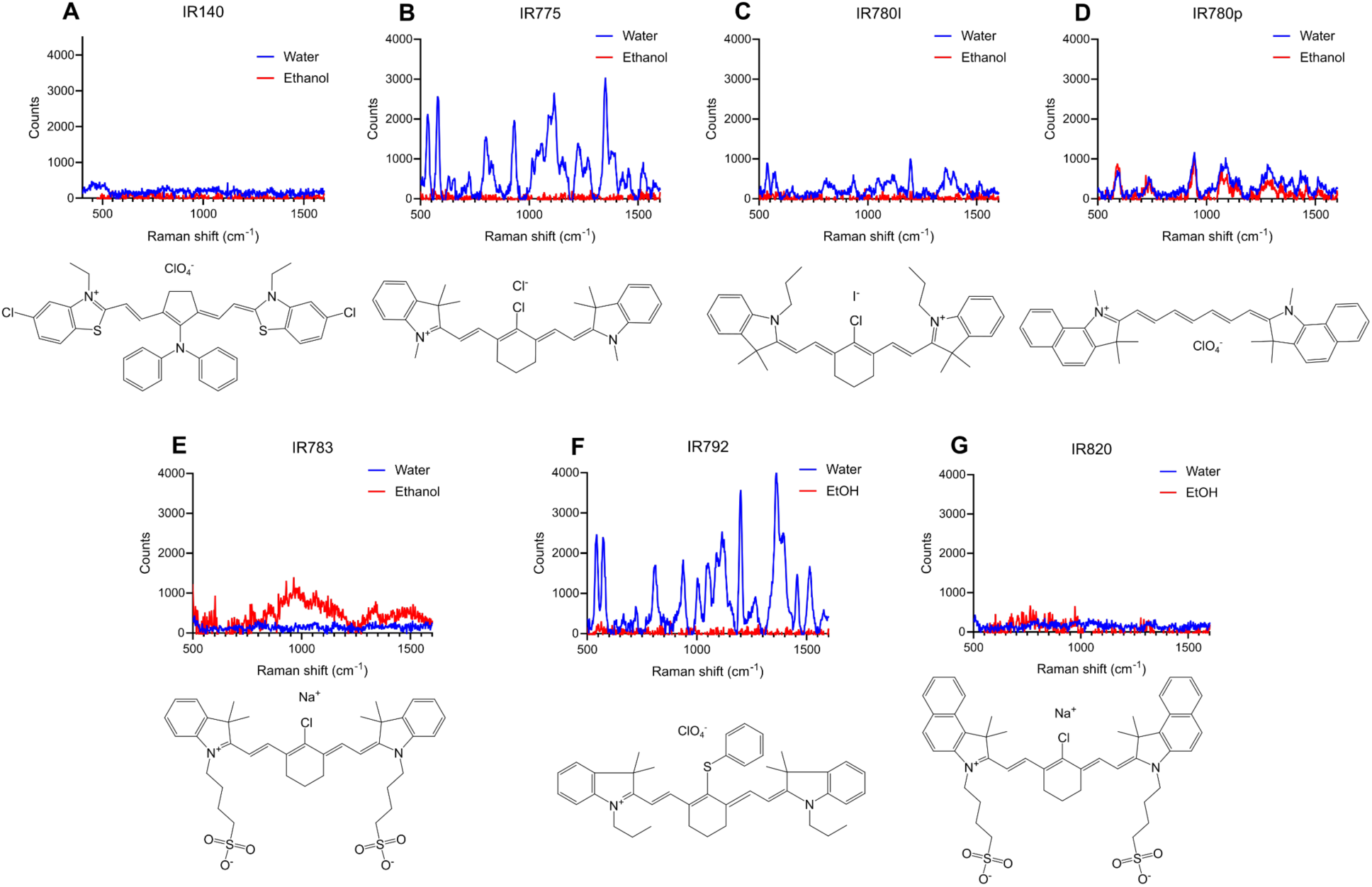
SERS spectra of AuNStars in water and ethanol immediately after the addition of each cyanine dye. (A) IR140, (B) IR775, (C) IR780I, (D) IR780p, (E) IR783, (F) IR792, and (G) IR820. Spectra were collected from AuNStar suspensions with an extinction of 0.3 at the LSPR. All spectra were baselined in MATLAB. All measurements were performed using a fiber optic Raman probe, 785 nm laser, 40 mW, 100 ms integration time, 1 accumulation. The EtOH and cuvette Raman background spectrum were removed by spectral subtraction.

Comparison of the SERRS responses revealed a clear charge- and structure-dependent relationship in signal intensity across the dyes (**Table 1**). Negatively charged IR dyes produced no detectable SERS in either water or ethanol, consistent with electrostatic repulsion between the negatively charged AuNStar surface (zeta potential = -46.8 ± 0.4 mV) and the anionic dye molecules. Conversely, positively charged dyes bearing a cyclohexene linker between indole moieties, specifically IR792, IR775, and IR780I, yielded measurable SERS signals when dispersed in water but not in ethanol. This suggests that adsorption of these dyes to the AuNStar surface occurred preferentially in aqueous media, whereas the more nonpolar ethanolic environment weakens the dye-surface interaction. Notably, IR780p was the only dye that generated SERS in both water and ethanol, indicating an affinity for the gold surface across a broader range of chemical environments.

**Table 1:** SERS response from AuNStars in water and ethanol in the presence of cyanine dyes.

| *IR dye* | *SERS in H<sub>2</sub>O* | *SERS in EtOH* | *Dye charge* |
| --- | --- | --- | --- |
| IR820 | X | X | - |
| IR783 | X | X | - |
| IR792 | ✓ | X | + |
| IR780p | ✓ | ✓ | + |
| IR780I | ✓ | NA | + |
| IR140 | X | X | + |
| IR775 | ✓ | X | + |

The Stöber process was used to encapsulate silica using a mixture of isopropanol, ethanol, and water in a 10:4:1 (v:v:v) ratio. Based on the results of **Table 1**, IR780p was selected as the Raman reporter most likely to adsorb efficiently to the AuNStar surface during silication and provide strong SERS signals post-encapsulation. The dye was introduced into the silication mixture at a final concentration of 50 µM (**Figure 2A**). Successful formation of the silica shell was confirmed by TEM (**Figure 2B**) and by a red shift in the LSPR (**Figure 2C and Figure 2D**). The resulting AuNStar-SiO_2_ exhibited a strong SERRS spectrum attributable to surface-bound IR780p molecules (**Figure 2E**).

**Figure 2.**
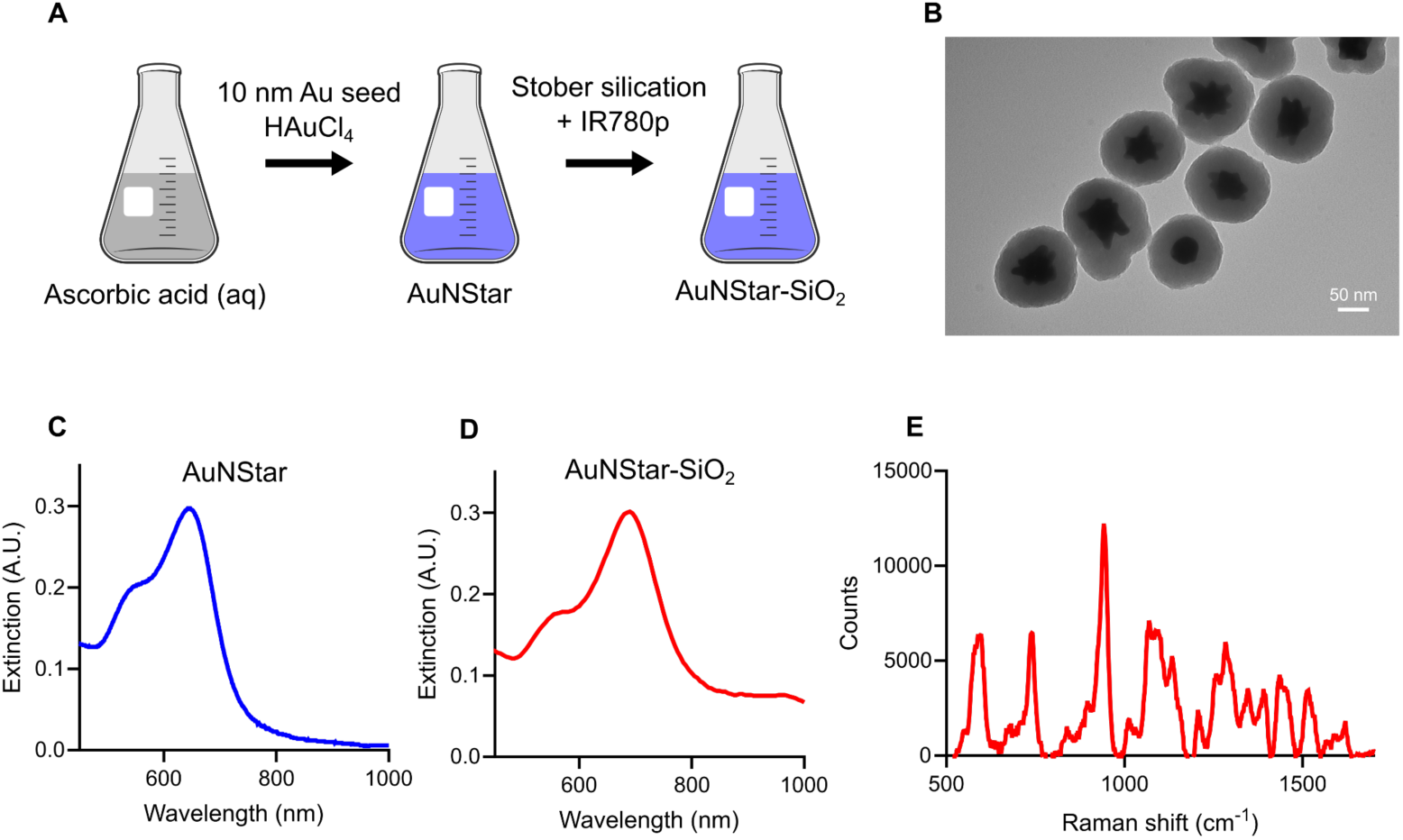
(A) Schematic of gold nanostar silication procedure. (B) TEM of silicated gold nanostars. (C) UV-vis spectrum of AuNStar suspension before silication. (D) UV-vis spectrum following silica encapsulation (AuNStar-SiO_2_). (E) SERRS spectrum of AuNStar-SiO_2_. All measurements were performed using a fiber optic Raman probe, 785 nm laser, 40 mW, 100 ms integration time, 1 accumulation. The cuvette Raman background spectrum was background subtracted.

### Size-separation of silicate AuNStar-SiO_2_

Following silication, a slight broadening of the AuNStar-SiO_2_ LSPR was observed (**Figure 2D**), consistent with previous reports on Stöber-based silica encapsulation of plasmonic AuNStars,^36,45^ and suggestive of some aggregation during the Stöber silication process and increased interparticle plasmonic coupling.^46^ Nanoparticle aggregation and the resulting nanogaps produce strong localized electromagnetic fields (hotspots) that can dominate the SERRS response.^27^ In contrast, individual AuNStars typically generate enhancement through strong electric fields localized at their sharp tips.^20^ To better understand the source of SERS enhancement in AuNStar-SiO_2_ samples, we employed continuous density gradient centrifugation to separate them by size and aggregation state.

AuNStar-SiO_2_ were separated in a continuous glycerol gradient (30-50%) over 20 minutes of centrifugation, and collected into seven fractions based on their relative positions in the column (**Figure 3A**). DLS revealed that hydrodynamic diameter increased with fraction number, indicating successful size-based separation (**Figure 3B**). Extinction spectra showed that the relative contribution of coupled plasmon modes, appearing as red-shifted bands beyond the ∼680 nm LSPR of individual AuNStar-SiO_2_, increased with higher fractions, confirming the enrichment of aggregates such as dimers, trimers, and oligomers (**Figure 3C**).^46^

**Figure 3:**
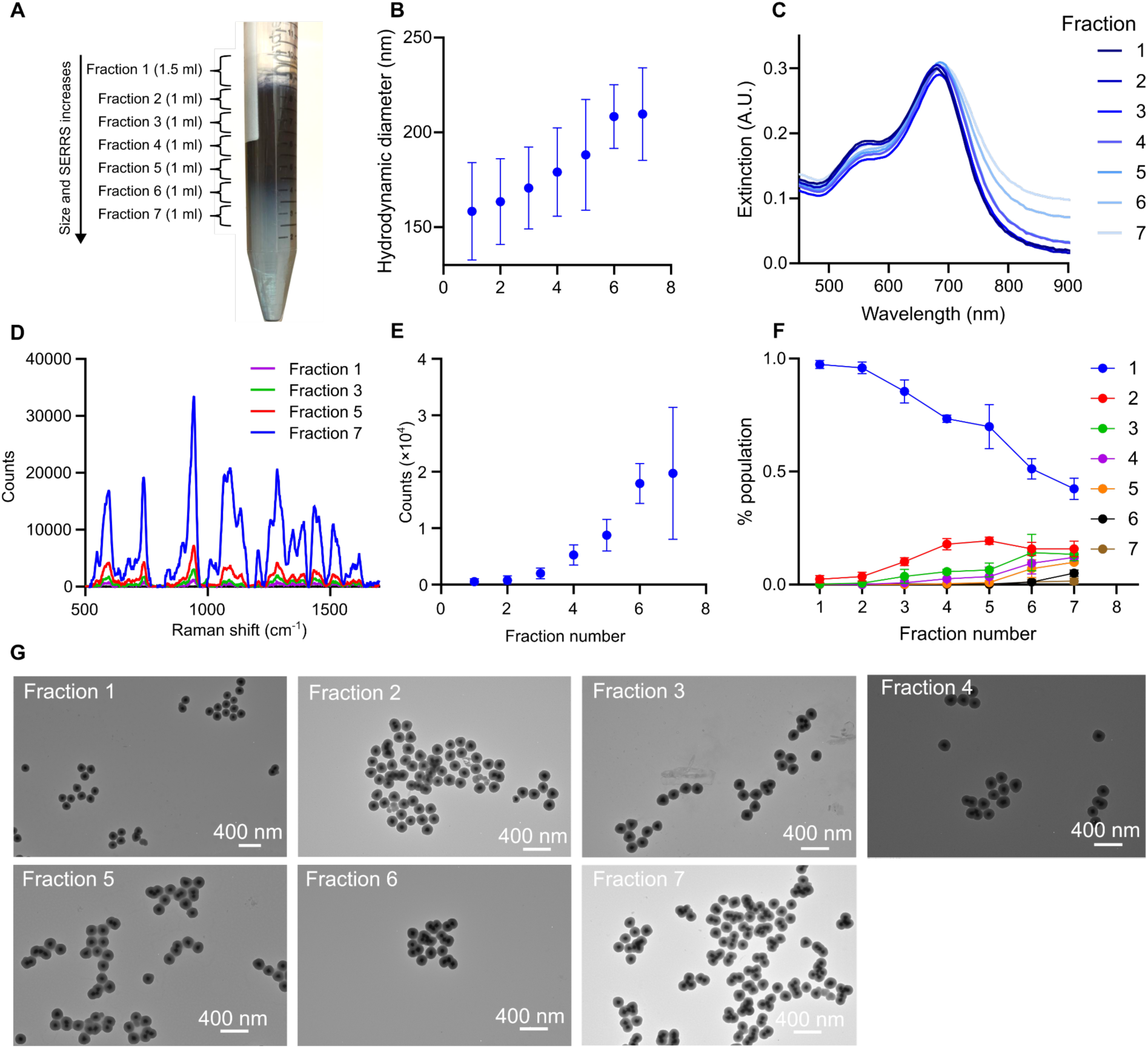
AuNStar-SiO_2_ fractionation and characterization. (A) Picture of 15 ml falcon tube after density gradient centrifugation of AuNStar-SiO_2_ sample. Fractions 1-7 indicate the volumes collected for further analysis. (B) Hydrodynamic diameter measurements of each fraction. Error bars represent the standard deviation of three AuNStar synthesis and purification experiments. (C) UV-vis extinction spectrum of fractions 1-7, an increase in extinction at wavelengths greater than the LSPR indicates aggregation and coupling between AuNStars within silica shells. (D) SERS spectra of fractions 1, 3, 5, and 7, collected from samples with an extinction of 0.3 at the LSPR. All measurements were performed using a fiber-optic Raman probe with a 785 nm, 40 mW laser, 100 ms integration time, and 1 accumulation. The cuvette Raman background spectrum was removed by spectral subtraction. (E) SERS intensity increases with fraction number and therefore size and interparticle coupling.(F) The percentage of AuNStar monomers, dimers, oligomers in each fraction counted from TEM images. >100 AuNStar-SiO_2_ and aggregates were counted for each fraction across three batches of AuNStar-SiO_2_. (G) Representative TEM images of Fractions 1-7.

SERRS intensity exhibited a pronounced dependence on the fraction number (**Figure 3D** and **Figure 3E**). Fractions 1 and 2, which contained the smallest nanoparticles, produced negligible SERRS signals, while spectral intensity increased progressively from fractions 3 to 7 as the hydrodynamic diameter also increased. TEM (**Figure 3F-G**) showed a clear correlation between aggregation and signal intensity: fraction 1 contained 97% monodisperse AuNStar-SiO_2_, whereas fraction 7 contained only 42% monomers, with the remaining particles forming silica-encapsulated multi-star aggregates (dimers-heptamers). Fraction 3 was selected for subsequent experiments as it consisted predominantly of monodisperse AuNStar-SiO_2_, exhibited moderately strong SERS activity, and showed no detectable higher-order aggregates comprising more than four AuNStar cores.

### Stability of AuNStar-SiO_2_ in physiological environments

During *in vitro* applications, AuNStar-SiO_2_ would be subjected to a range of pH environments, which may affect the stability of the silica shell and the interaction between the dye molecules and the nanostar surface. The extracellular environment of standard cell culture systems typically maintains a pH between 7.2 and 7.6. However, values as low as pH 6.4 have been reported in more advanced organoid and air-liquid interface models,^47^ and pHs above physiological levels can occur in cell culture medium under low CO_2_ pressure conditions.^48^ Furthermore, *in vivo* the tumor microenvironment can be acidified to pH ∼6.4 by the Warburg effect.^49^ During endocytosis, nanoparticles are generally trafficked to the lysosome via progressively more acidic compartments, reaching a pH ∼4 within the lysosome.^15^ We evaluated the structural and optical stability of our purified fraction 3 AuNStar-SiO_2_ nanoparticles in biologically relevant pH conditions by incubating them in phosphate-buffered saline (PBS) adjusted to pH 4, 6.5, 7.4, and 9. These conditions were designed to model lysosomal acidity, the near-neutral environment of culture media, and the alkaline conditions of basified high-bicarbonate cell culture medium at atmospheric CO_2_.

Fraction 3 AuNStar-SiO_2_ suspensions were monitored for 25 h in each pH using UV-vis spectroscopy (**Figure 4A-D**), SERS (**Figure 4E-H**), and DLS (**Figure 4I-L**). At pH 4, the SERRS intensity (**Figure 4A**) decreased gradually to approximately 60% of its initial value over 25 h. The LSPR blue-shifted by ∼100 nm (**Figure 4E**), consistent with morphological reshaping of the AuNStars into more spherical particles, as confirmed by TEM (**Figure S1**). The hydrodynamic diameter increased markedly, promoting nanoparticle sedimentation (**Figure 4I**), and the silica shell remained intact (**Figure 4M**). At pH 9, a two-fold increase in SERRS intensity was observed after 3 h (**Figure 4D**) and paired with an LSPR blue shifted by ∼20 nm (**Figure 4H**), indicating a reduction in the local dielectric environment from the loss of the silica shell. During this time, the DLS measurements showed a decrease in hydrodynamic diameter to ∼60 nm (**Figure 4L**), which is consistent with the starting size of bare AuNStars. TEM at 25 h showed bare AuNStars without silica shells (**Figure 4P**), demonstrating that at high pH the silica shell is unstable. At pH 6.5, the AuNStar-SiO_2_ nanoparticles exhibited intermediate stability. During the first 12 h of incubation, the localized surface plasmon resonance (LSPR) underwent a ∼100 nm blue shift (**Figure 4F**). Prolonged incubation (12-25 h) resulted in complete hydrolysis of the silica shell (**Figure 4N**), releasing the AuNStars into solution. The liberated nanostars subsequently aggregated and sedimented, as evidenced by the high variability in SERRS intensity between replicates (**Figure 4B**), broadening of the plasmon band (**Figure 4F**), and the absence of an intact silica shell in TEM images at 25 h (**Figure 4N**). At pH 7.4, intermediate behavior was also observed. The silica shell was hydrolyzed more slowly, resulting in the release of AuNStars, which then aggregated (**Figure 4G**). The SERRS signal initially decreased, then increased threefold at 4 h (**Figure 4C**). We attribute this signal increase to interparticle plasmon coupling. Over the subsequent 20 h, this enhancement decayed as the aggregates irreversibly sedimented. UV-vis spectra showed an initial ∼20 nm blue shift within 2 h, reflecting the reduced surface dielectric from silica loss, followed by spectral broadening associated with interparticle coupling (**Figure 4G**). Hydrodynamic diameter analysis corroborated this, with an initial decrease followed by growth at 4 h (**Figure S2**). TEM images confirmed silica shell hydrolysis and exposure of bare AuNStars (**Figure 4O**). In contrast, AuNStar-SiO_2_ dispersed in ultrapure water exhibited no significant changes in SERRS intensity, optical properties, or morphology over the 25 h observation period (**Figure S3**).

**Figure 4.**
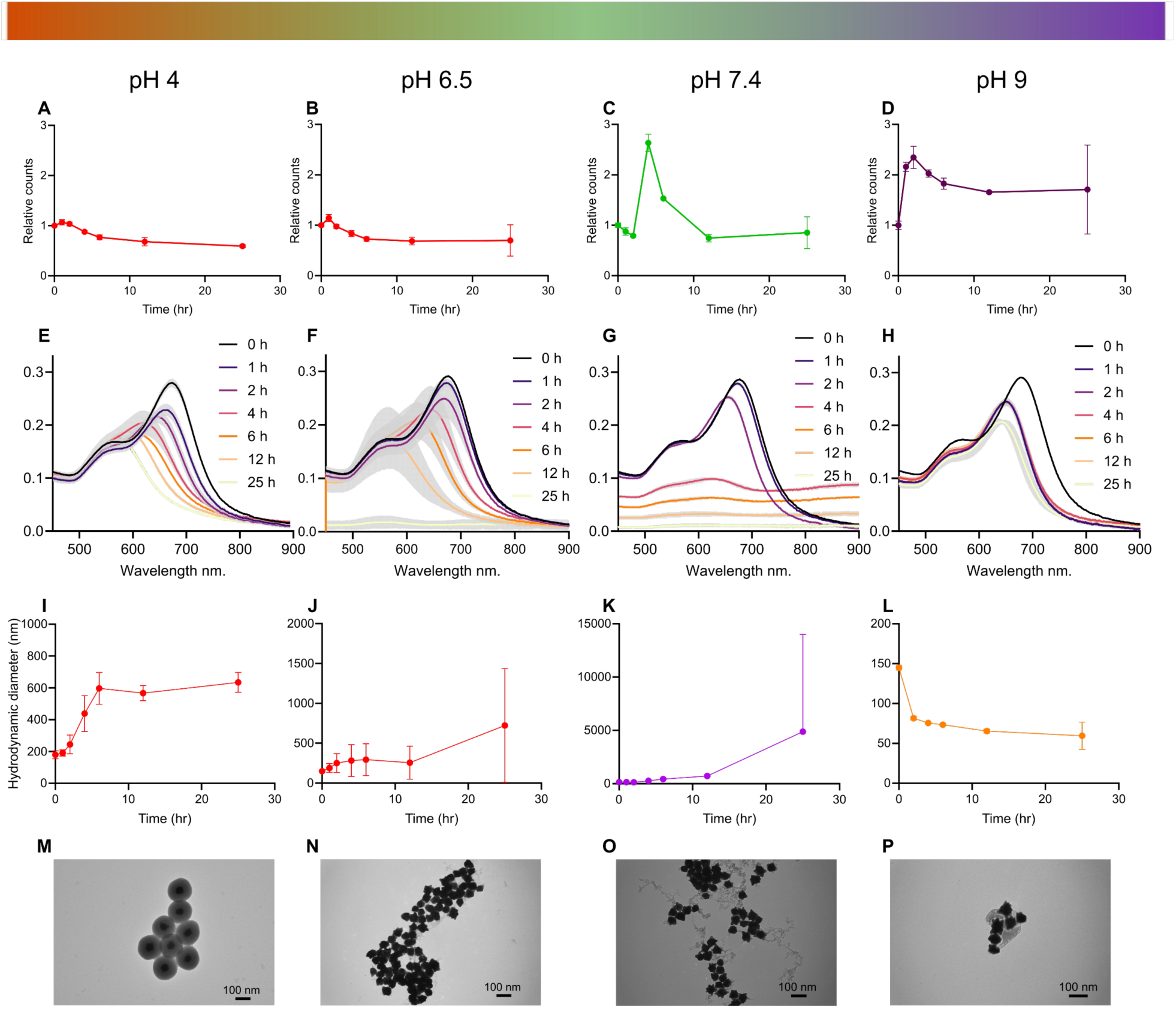
Stability of AuNStar-SiO2 nanoparticles in PBS adjusted to pH 4, 6.5, 7.4, and 9 during 25 h incubation at 37 °C. The optical and physical properties of AuNStar-SiO2 were measured at defined time points throughout the incubation. All samples were vortexed prior to analysis. The SERRS intensity of the band at 940 cm⁻¹ was recorded at each time point for (A) pH 4, (B) pH 6.5, (C) pH 7.4, and (D) pH 9. All measurements were performed using a fiber-optic Raman probe with a 785 nm, 40 mW laser, 100 ms integration time, and 1 accumulation. Error bars represent the standard deviation from n = 3 technical replicates. The UV-vis extinction spectra of AuNStar-SiO2 acquired in (E) pH 4, (F) pH 6.5, (G) pH 7.4, and (H) pH 9. The shaded regions denote the standard deviation (n = 3). Nanoparticle size was measured with DLS for each point at (I) pH 4, (J) pH 6.5, (K) pH 7.4, and (L) pH 9, (n = 3). The starting sizes for pH 4 and pH 9 differ because of the rapid rate of aggregation and silica hydrolysis, respectively. TEM images of AuNStar-SiO2 following 25 h incubation at (M) pH 4, (N) pH 6.5, (O) pH 7.4, and (P) pH 9.

### *In vitro* AuNStar-SiO_2_ hydrolysis and imaging

Having demonstrated that pH-dependent silica hydrolysis modulates the SERRS intensity of AuNStar-SiO_2_ contrast agents, we examined whether similar effects occur in biologically relevant *in vitro* environments when nanoparticles are endocytosed by cells. We investigated two conditions: near-neutral extracellular pH 7.4, typical of *in vitro* cell culture, and an acidic extracellular environment of pH 6.4 mimicking tumor extracellular acidification.^49^ By varying extracellular pH and quantifying intracellular SERRS responses, we aimed to establish how local acidity impacts nanoparticle stability and SERRS signal output, providing insight into the performance and reliability of silica-coated SERRS probes under conditions relevant to tumor biology.

SW48 cells were incubated with AuNStar-SiO_2_ for 4 h at pH 6.4, and at pH 7.4, (**Figure 5A**). The 4 h time point was selected to allow nanoparticle endocytosis^50^ and to permit complete silica dissolution at pH 7.4 but only partial dissolution at pH 6.4 (**Figure 4F-G**). Raman maps of the cells were analyzed pixel-wise to quantify SERRS differences between conditions (**Figure 5B**). Pixels with a signal-to-noise ratio >3 at the 940 cm^-1^ IR780p mode were defined as SERRS-positive, while all others were thresholded to zero. SW48 cells cultured at pH 6.4 exhibited nearly a threefold higher SERRS-positive pixel density per µm² of cell area compared with cells incubated at pH 7.4 (**Figure 5C**).

**Figure 5:**
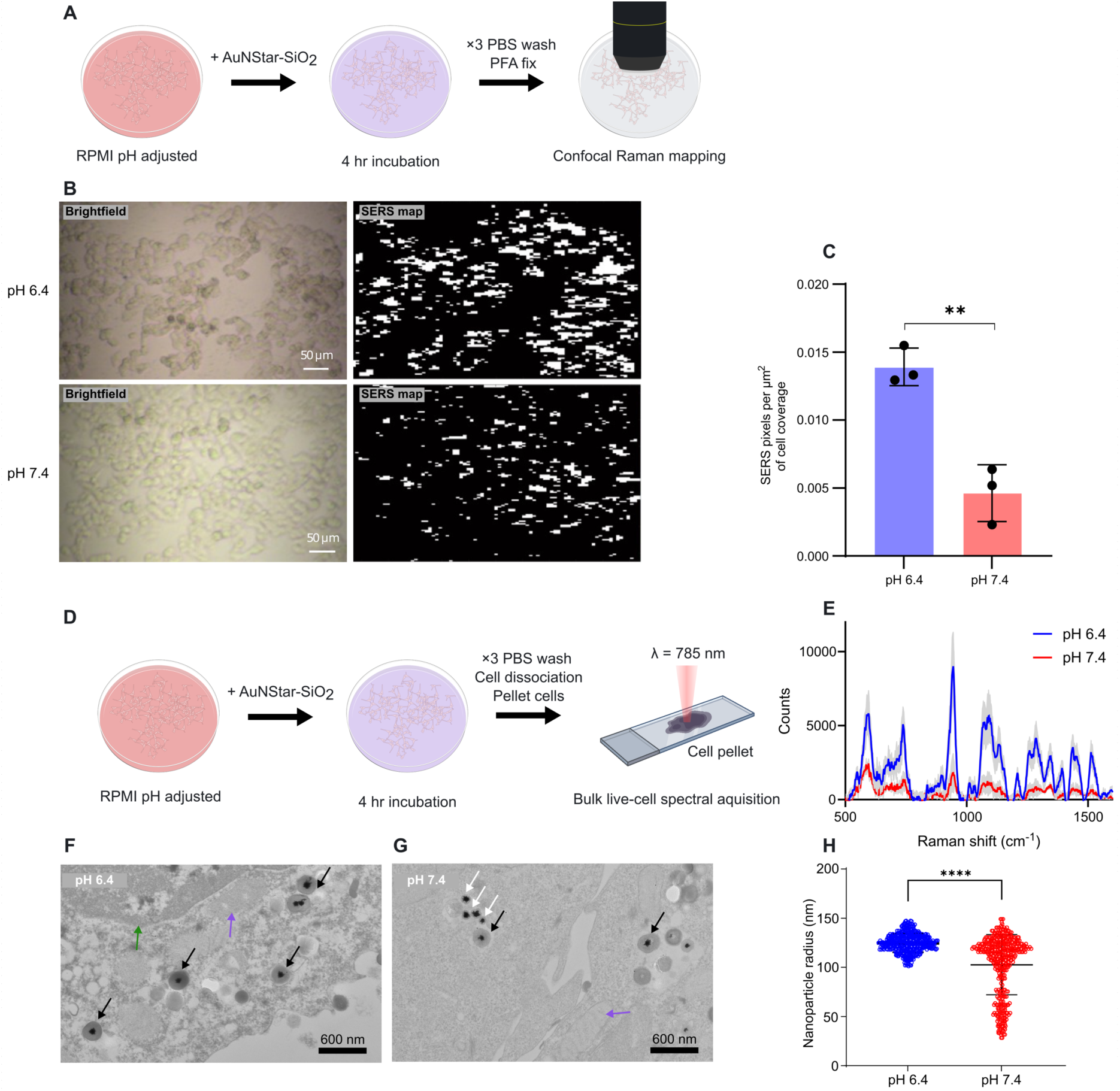
AuNStar-SiO_2_ as *in vitro* imaging agents under physiological and acidic conditions. (A) Schematic of cell mapping experiment. SW48 cells were incubated with AuNStar-SiO_2_ in regular medium (pH 7.4) and acidic medium (pH 6.4) for 4 hours before fixing and mapping with a Raman confocal microscope. (B) Brightfield and Raman maps of fixed cells after incubation with AuNStar-SiO_2_ at pH 6.4 and pH 7.4. Raman maps were created by masking the cell area in the brightfield and assigning pixels a binary value of 1 or 0 based on whether the spectral intensity at 940 cm⁻¹ exceeded 3× the noise level. Maps were collected with a 10x objective, 10% laser power, 0.1 s acquisition. (C) The density of pixels with SERRS intensity > 3× the noise level per µm^2^ of cell coverage for cells incubated at pH 6.4 and pH 7.4. Data points are from independent biological replicates. Each data point is the mean of 3 maps from a single sample (t-test, p = 0.0049). (D) Schematic of SERRS analysis of live-cell pellet following 4 h of incubation with AuNStar-SiO_2_ at each pH. (E) SERRS spectra of pelleted SW48 cells following incubation at each pH. The shaded area represents the standard deviation of n=3 biological repeats. Spectra were collected with a Raman probe, 785 nm laser, 1 ms exposure time, single acquisition, and a laser power of 40 mW. (F) TEM image of intracellular AuNStar-SiO_2_ after 4 h of incubation at pH 6.4. Black arrows highlight nanoparticles with a complete silica shell, and white arrows highlight nanoparticles that have undergone silica hydrolysis. The green arrow highlights the cell nucleus, and the purple arrow highlights a mitochondrion. (G) TEM image of intracellular AuNStar-SiO_2_ after 4 h of incubation at pH 7.4. (H) Intracellular nanoparticle size in SW48 cells incubated with nanoparticles for 4 hours at pH 6.4 and pH 7.4. (p < 0.0001).

To validate these findings, we repeated nanoparticle incubation at pH 6.4 and 7.4, washed and pelleted the cells, and acquired bulk SERRS spectra from the live-cell pellet deposited on a glass slide using a fiber-optic Raman probe (**Figure 5D-E**). Consistent with the cell-mapping data, cells exposed to AuNStar-SiO_2_ at pH 6.4 exhibited a statistically greater SERRS intensity than cells treated at pH 7.4, by a factor of 5 this case (**Figure S4**). TEM images of intracellular AuNStar-SiO_2_ revealed nanoparticles internalized by SW48 cells at pH 6.4 had intact silica shells with little size heterogeneity (**Figure 5F**). At pH 7.4, AuNStar-SiO_2_ had a range of silica shell thicknesses between complete shell and complete dissolution (**Figure 5G**). The statistically significant decrease in intracellular nanoparticle size between pH 6.4 and pH 7.4 indicated a greater rate of silica shell hydrolysis at pH 7.4 prior to endocytosis (**Figure 5H**).

## DISCUSSION

In this study, we characterized SERRS AuNStar-SiO_2_ for *in vitro* imaging applications and elucidated the factors that affect signal and nanoparticle stability in biological environments. We first selected the optimal dye molecule for AuNStars SERRS by examining the effects of dye charge and solvent on spectral intensity. We observed that IR dyes published in the literature as SERRS reporters exhibited varying affinities for the AuNStar nanoparticle surface. In an aqueous environment, coulombic interactions were the primary factor. Our AuNStars have a zeta potential of -46.8±0.4 mV, and positive IR dyes, IR792, IR780p, IR780I and IR775 all showed SERS, but no signal was visible from negatively charged dyes such as IR820 and IR783 (**Figure 1A-G**). Although IR140 is a positively charged molecule, it exhibited poor water solubility, which prevented SERRS acquisition. Surprisingly, only IR780p produced SERRS when added in ethanol. While IR792 showed the strongest signal in water, it generated no SERS in ethanol; therefore, IR780p was selected for its surface affinity in ethanol, which we reasoned would increase the probability of dye molecules adhering to the AuNStars during the Stöber silication process.

Following silication with dye molecule IR780p, AuNStar-SiO_2_ produced strong SERRS signal (**Figure 2E**), however size separation of nanoparticle subpopulations using density gradient centrifugation showed the majority of SERRS was produced by coupled-AuNStar aggregates encapsulated in silica shells. The majority of monomeric fractions 1 and 2 showed very low SERS intensity (**Figure 3E-G**), suggesting that, once encapsulated, the NIR dye IR780p was not located within the intense electric field regions at the nanostar tips. These observations suggested that IR780p molecules are unable to adsorb directly onto the gold surface at the interface with silica at the star tip. Dye molecules trapped within the silica shell are either too sparse or too distant from the plasmonic surface to experience significant electromagnetic enhancement. Instead, the strong SERRS observed in aggregated fractions appeared to originate from IR780p molecules confined within interparticle junctions at Au-Au interfaces that act as hotspots. These results highlight that the origin of SERRS enhancement in AuNStar systems can shift following silication. Our bare AuNStars generated SERRS when dye molecules were added, indicating that the electric fields at their spike tips were sufficient for enhancement. However, the majority of SERRS signals in their silicated counterparts was generated by coupled AuNStars within silica shells, demonstrating that even in star-based systems, interparticle gaps can dominate SERS signal. Fraction 3 was selected for stability and *in vitro* studies because it had a high monomer content (85%), stronger SERRS than fractions 1 and 2, and no detected higher-order aggregates of 4+.

We explored the stability of AuNStar-SiO_2_ fraction 3 across a series of increasingly acidic environments to simulate the pH changes that occur during nanoparticle endocytosis. The stability of silica nanoparticles has been investigated primarily in the context of drug-loaded silica systems, where the degradation rate determines the location and timing of nanoparticle payload release.^51–53^ These studies show that silica hydrolysis is most pronounced for particles with low condensation silica networks and high surface-to-volume ratios, particularly under alkaline, high-salt conditions. To the best of our knowledge, the role of silica nanoshell stability in plasmonic applications has received limited attention. The importance of characterizing silica shell stability extends beyond the biomedical applications discussed in this paper, as silica-coated SE(R)RS nanoparticles are used in anti-counterfeiting barcodes and security labels,^54–56^ food safety,^57–59^ and environmental monitoring^60,61^ where stability against pH-driven hydrolytic weathering is vital. In our work, the silica nanoshells in AuNStar-SiO_2_ underwent hydrolytic degradation under physiological and alkaline conditions, thereby releasing the plasmonic AuNStar core and changing the nanoparticles optical properties and surface-dye interactions. We observed the most pronounced optical effects at pH 7.4 (**Figure 4C**) where AuNStars were released within four hours and rapidly aggregated, temporarily increasing the SERRS signal before sedimentation reduced the signal. At pH 6.5, the shell hydrolysis rate was lower, and complete silica shell dissolution and AuNStar aggregation were only observed at 25 h (**Figure 4F**). At pH 4, the silica shell remained intact after 25 h of incubation, indicating reduced silica hydrolysis at low acidic pH. However, a redshift of 100 nm was observed in the AuNStar-SiO_2_ LSPR, which we confirmed via TEM was a result of AuNStars within the silica shell becoming more spherical at pH 4 (**Figure S1**). These results highlight the physicochemical changes that silica-encapsulated plasmonic nanoparticles can undergo in environments representative of in-vitro cell culture. In sensing applications where signal intensity or spectral features are used to quantify nanoparticle internalization or targeting efficiency, degradation of the silica shell and the associated changes in optical properties could significantly influence the assay outcomes. These results motivated us to explore silica shell degradation *in vitro* following endocytosis and to characterize the resulting impact on SERRS intensity from intracellular AuNStar-SiO_2_.

Two pH conditions were tested for AuNStar-SiO_2_ endocytosis by SW48 cells. pH 7.4 was chosen as the standard pH for RPMI buffered with 5% CO_2_, a condition widely used in cell culture. In this experiment, the silica shell of nanoparticles was hydrolyzed in the extracellular space, and this degradation slowed once the nanoparticles were endocytosed and the pH of their environment lowered. TEM images of intracellular AuNStar-SiO_2_ nanoparticles endocytosed at pH 7.4 showed a range of nanoparticle sizes (**Figure 5H**), reflecting the different extents of silica hydrolysis at the time of internalization. The results of **Figure 4G** indicate that complete silica hydrolysis at pH 7.4 occurs in approximately 4 hours. Endocytosis is a rapid process on the seconds-minutes timescale,^50^ which allows cells to endocytose AuNStar-SiO_2_ with complete shells at 0 h and fully degraded shells at 4 h. This progressive decrease in pH effectively arrested the hydrolysis of the nanoparticle silica shell at the point of endocytosis. The second condition we explored was RPMI at pH 6.4. In this experiment, limited silica hydrolysis occurred in the extracellular environment during the 4 h endocytosis period and internalized AuNStar-SiO_2_ had a more homogeneous size. Unlike the short-lived threefold signal enhancement observed during silica dissolution in suspension (**Figure 4C**), silica hydrolysis appears to reduce SERRS signal when nanoparticles are endocytosed (**Figure 5C and 5E**). These two seemingly conflicting results can be rationalized by the same effect: the hydrolysis of the silica shell and subsequent release of the Raman dye molecules from the silica matrix and interparticle coupling. In a suspension, silica-shell dissolution temporarily increased the signal when loss of silica shells drove electrostatic aggregation and created interparticle hotspots, which increased SERRS before complete sedimentation reduced SERRS (**Figure 4C**). When the silica shell is hydrolyzed and the nanoparticles are endocytosed, SERRS intensity decreases, most likely because competing nanoscale and microscale effects prevent interparticle coupling of dye molecules at the hotspot. At the nanoscale, protein binding to the surface of nanostars could also prevent the localization of dye molecules in interparticle cavities, while encapsulation in separate endocytic vesicles keeps nanostars separated at the microscale and prevents plasmonic coupling.

Silica shells are used in many *in vitro* and *in vivo* SERRS imaging platforms. In most applications a polyethylene glycol (PEG) layer is added to the silica shell to create a hydrophilic barrier to reduce nonspecific protein adsorption, limit aggregation, and improve circulation time *in vivo*.^8,11,36,45^ In literature examining payload release from silica nanoshells, PEG has been shown to reduce water access to the silica shell and slow degradation kinetcs.^53^ However, an increasing body of research is also highlighting PEG-specific immune responses such as complement activation-related pseudoallergy (CARPA),^62–64^ and accelerated blood clearance, driven by anti-PEG antibodies developed following the first treatment and reducing treatment efficacy in subsequent doses.^65^ In this study, we intentionally examined the silicated nanoparticles in the absence of PEG to isolate the intrinsic stability of the silica shell in biologically relevant environments in the absence of a surface coating that induces specific biological responses and may in the future be surpassed by less immunogenic coatings. Our findings are important because silica encapsulation layers are often implicitly assumed to be chemically stable and inert under physiological conditions. By decoupling silica stability from the protective effects of PEG, we directly probe how mild biological environments influence silica shell integrity, and in turn, SERRS spectral behavior. Our results demonstrate that commonly used silica shells are highly susceptible to chemical degradation even under modest pH changes representative of biological milieus. Importantly, silica hydrolysis was found to have divergent effects on SERRS performance, capable of either enhancing or suppressing signal intensity depending on the local chemical environment and measurement time point. Our experiments in cuvettes using pH-adjust PBS provided key mechanistic insights, but ultimately, we observed a decrease in spectral intensity, rather than the expected increase, when the same process of AuNStar-SiO_2_ silica degradation occurred *in vitro*. These findings reveal silica degradation as a previously underappreciated source of spectral variability in SERRS nanoprobes and highlight the need for multimodal in situ characterization of biologically sensitive nanoparticles across the range of conditions encountered in cells, tissues, and organs to fully characterize nano-bio interactions and accurately interpret spectral output.

Achieving reliable SERRS-based quantification of nanoparticle accumulation in biological tissues has long been a goal of the field. Such quantitative applications require nanoprobes that maintain structural integrity and produce reproducible signal intensities in complex biological environments. Our data identify silica hydrolysis as a critical factor that can undermine this reproducibility, showing that relatively small pH variations, well within those encountered during biological trafficking, can significantly alter SERRS imaging performance. At the same time, these results suggest that degradation of the silica shell should not be viewed solely as a limitation. Instead, environment-responsive silica behavior may be intentionally leveraged to modulate SERRS enhancement, offering new opportunities to engineer nanoprobes whose spectral output is tuned by local biochemical conditions. Furthermore, in situ nanoparticle size reduction via silica hydrolysis may be a promising strategy to prevent long-term sequestration of plasmonic nanoparticles in the reticuloendothelial system by shrinking nanoparticle size below the threshold for renal clearance,^66–68^ thus preventing potential changes in gene expression induced by chronic gold nanoparticle exposure and increasing translational potential.^69^ Together, these findings highlight the need to reconsider assumptions of silica inertness in SERRS probe design and motivate future strategies that balance stability, sensitivity, and environmental responsiveness for quantitative molecular imaging.

## CONCLUSION

In this work, we explored the mechanistic drivers behind SERRS signal generation by AuNStar-SiO_2_ in suspensions and in cellular environments. We demonstrated a relationship between SERRS signal intensity and interparticle plasmonic coupling by using density gradient centrifugation to separate AuNStar-SiO_2_ subpopulations into size-dependent fractions. Interparticle coupling in AuNStar-SiO_2_ was shown to be the main driver of SERRS intensity in our nanoparticle system. Further, and more importantly, we showed that silica encapsulation alone may not be a pH-stable protection strategy for nanoparticles in the neutral-alkaline environments of *in vitro* cell culture. Complete dissolution of silica nanoshells temporarily induced a significant increase in the SERRS signal when experiments were performed in PBS. This led us to explore silica-shell hydrolysis *in vitro* after AuNStar-SiO_2_ endocytosis and to show that, unlike in PBS, the SERRS signal is suppressed following shell loss. We used TEM to confirm increased rates of silica shell hydrolysis in cells exposed to nanoparticles in pH 7.4 cell culture medium than those exposed in mildly acidified medium. We concluded that when nanoparticles are free to aggregate and the dye molecule can diffuse into associated hotspots, such as in PBS solution, a short-lived SERRS increase follows silica shell dissolution. However, in the intracellular space, silica dissolution dampens SERRS signal because hotspots containing dye molecules are less likely to form, and dye diffusion to the hotspots is hindered by competing biomolecules. Together, these results demonstrate that SERRS performance in biologically relevant systems is governed by subpopulation heterogeneity, environment degradation, and cellular trafficking, revealing multiple, compounding sources of variability. Our findings also highlight an opportunity to fine-tune silica chemistry and create a new generation of SERRS probes that achieve greater signal strength, payload delivery, and excretion by responding to environmental cues.

## ACKNOWLEDGMENTS

This work was supported by the following grants: F.N. gratefully acknowledges support from the following awards: NIH/NCI R00CA266921, Claudia Adams Barr Award for Innovative Cancer Research, Optica Foundation Challenge Award, Friends of DFCI, and DFCI start-up funds. The authors would like to thank Kyle Smith at the Beth Israel Deaconess Medical Center Electron Microscopy Core for fixing, staining, and sectioning the cells for TEM.

## AUTHOR CONTRIBUTIONS

W.H.S. designed and conducted experiments, analyzed the data, and wrote and edited the manuscript. S.P. conducted experiments, analyzed the data, and edited the manuscript. F.N. designed experiments, analyzed the data, funded and supervised the studies, and wrote the manuscript. All authors reviewed the manuscript prior to submission.

