## Supporting Information for "pH-Dependent Silica Nanoshell Degradation Influences SERRS Enhancement in Biological Environments"

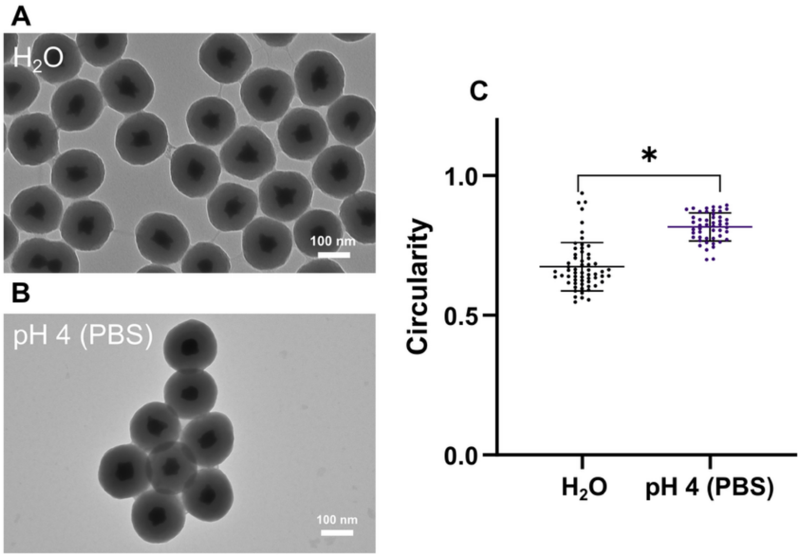


**Figure S1: reshaping of silicate gold nanostars within silica shell at pH 4. (A) TEM image of AuNStar-$SiO_2$ stored in water for 24 hours. (B) TEM image of AuNSTar-$SiO_2$ stored in PBS at pH 4 for 24 hours. (C) Circularity of AuNStars stored in water and pH 4 PBS for 24 hours. Circularity was measured in ImageJ by highlighting the AuNStar core and using the equation $C = \frac{4\pi A}{p^2}$ , where *C* is circularity, *A* is the area of the AuNStar and *P* is the perimeter of the AuNStar.**

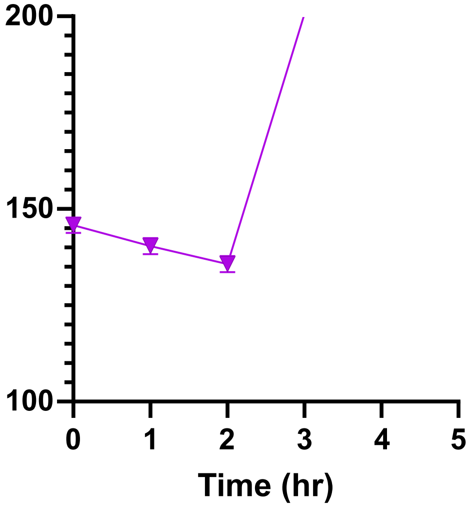


**Figure S2: DLS measurements of AuNStar-SiO2 during the first 2 hours of incubation in PBS at pH 7.4. The hydrodynamic diameter initially decreases as the silica shell is hydrolyzed. Nanoparticle size subsequently increases as the released AuNStars aggregate.**

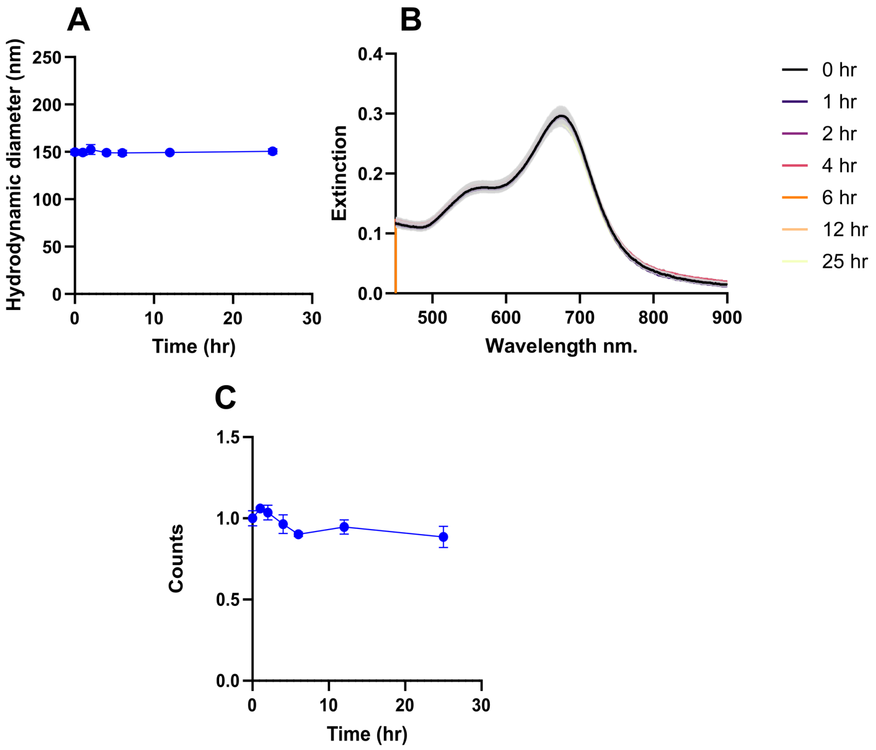


**Figure S3: Stability of AuNStars in ultrapure water over 25 hrs, (A) hydrodynamic diameter measured by DLS, (B) UV-vis spectra and (C) SERS spectra. Three technical repeats were performed per time point.**

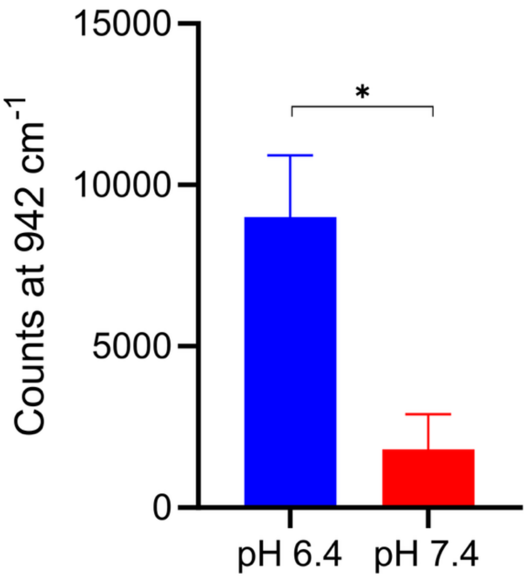


**Figure S4: Intensity of SERS peak at 942 cm-1 from cells incubated with AuNStarSiO2 for 4 hours at pH 6.4 and pH 7.4 (p = 0.02, n=3).**
